# Pan-cancer mutational landscape of the PPAR pathway reveals universal patterns of dysregulated metabolism and interactions with tumor immunity and hypoxia

**DOI:** 10.1101/563676

**Authors:** Wai Hoong Chang, Alvina G. Lai

## Abstract

Peroxisome proliferator-activated receptors (PPARs) are a family of nuclear receptors that regulate lipid metabolism and bioenergetic demands within living systems. Consequently, aberrant expression of PPAR genes could predispose individuals to diseases including cancer. PPAR signaling exerts pleiotropic functions in cancer, yet, little is known about the interactions between genetic and transcriptional events of pathway genes in a pan-cancer context. Employing multidimensional datasets of over 18,000 patients involving 21 cancers, we performed systematic characterization on copy number alteration and differential transcript expression of 74 PPAR pathway genes. We identified 18 putative driver candidates demonstrating mutually exclusive patterns of loss- and gain-of-function phenotypes. These driver genes successfully predicted patient survival rates in bladder, renal, glioma, liver and stomach/esophageal cancers. Dysregulated PPAR signaling in these cancers converged on common downstream pathways associated with multiple metabolic processes. Moreover, clinically-relevant relationships between PPARs and hypoxia were observed where hypoxia further aggravates disease phenotypes in tumor subtypes with aberrant PPAR signaling. In glioma samples, including astrocytoma and oligoastrocytoma, PPAR hyperactivation is associated with immunosuppression through increased regulatory T cell expression. Our analysis reveals underappreciated levels of diversity and conservation in PPAR genes that could lay the groundwork for therapeutic strategies targeting tumor metabolism, immunity and hypoxia.

## Introduction

A series of genetic and phenotypic changes must take place during malignant transformation to rewire cellular programs and signal transduction pathways into a permissive state that facilitates tumor survival(1–4). Metabolic reprogramming, which occurs through the attainment of genetic and epigenetic mutations, is often a critical step in tumorigenesis because cancer cells have a unique requirement for increase glucose uptake and lactate fermentation even in the presence of oxygen(5, 6). Tumor cells also have increased fatty acid oxidation and turnover(7), and this is perhaps not unforeseen because fatty acids yield twice as much energy as glucose. Tumor cells with increased de novo fatty acid synthesis through the upregulation of fatty acid synthase are also often more aggressive(8). Fatty acid oxidation is activated by upstream effectors involving signal transduction pathways such as peroxisome proliferator-activated receptor (PPAR) and AMP-activated protein kinase(5, 9). Given their bioenergetic dependence on fatty acid oxidation, a better understanding of how lipid pathways are dysregulated in cancer cells will be crucial for successful therapy.

PPARs are a family of nuclear receptor transcription factors involved in regulating metabolic homeostasis whose activity is regulated by fatty acid ligands(10). Three categories of PPARs have been identified in humans; PPARα, PPARβ/δ, and PPARγ. Once activated, PPARs heterodimerize with retinoid X receptors (RXRs) to regulate the expression of downstream target genes harboring peroxisome proliferator response element motifs. Given the availability of PPAR agonists and antagonists, investigations on the role of PPAR signaling in cancer has begun to gain momentum. PPAR agonists may promote or suppress tumor formation depending on the cellular context. PPARγ agonists such as pioglitazone and troglitazone are effective inhibitors of cell growth in colorectal(11–13), melanoma(14) and ovarian cancers(15). PPARγ agonists could also inhibit the proliferation and induce cell cycle arrest in brain tumor stem cells in a dose-dependent manner(16). In contrast, treatment with a PPARγ antagonist depletes tumorsphere formation in ERBB2+ breast cancer stem cells, suggesting that PPARγ is essential for breast cancer stem cell maintenance(17). PPARδ is also required for hematopoietic stem cell maintenance, and its inhibition prevents asymmetrical cell division required for stem cell self-renewal(18).

The ubiquitous impact of PPAR signaling across diverse cancer types necessitates a systematic curation of genomic and transcriptomic alterations of all PPAR pathway genes using a pan-cancer approach, which may offer unprecedented insights into elucidating novel druggable targets. We performed an integrated analysis on PPAR pathway genes using genomic, transcriptomic and clinical data from 18,484 patients involving 21 cancer types (Fig. 1A). We discovered common and distinct patterns of genetic mutations and dysregulated transcript expression converging on a core set of 18 putative driver genes that harbor clinically-relevant prognostic information in multiple cancer types. Given its pervasive influence in numerous biological processes, we also examined the crosstalk between PPAR signaling and tumor hypoxia or tumor immunity. Oncogenic variants of the PPAR pathway identified in this study can be integrated into current initiatives on personalized therapy employing PPAR agonists to target fatty acid metabolism in conjunction with first-line treatments.

## Methods

### Gene set and cancer cohorts

Seventy-four PPAR pathway genes were retrieved from the Kyoto Encyclopedia of Genes and Genomes (KEGG) database listed in Table S1. Transcriptomic, genomic and clinical datasets of 21 cancer types generated by The Cancer Genome Atlas (TCGA) were retrieved from the Broad Institute GDAC Firehose website(19).

### Somatic copy number alterations analyses

GISTIC Level 4 copy number variation profiles for each cancer type were downloaded from Firehose. To determine gene amplifications and deletions, we utilized GISTIC gene-level tables that provided discrete amplification and deletion indicators(20). Samples with GISTIC values higher than the maximum copy-ratio for each chromosome arm (> +2) were annotated as ‘deep amplification’ events. In contrast, samples with values lower than the minimum copy-ratio for each chromosome arm (< −2) were annotated as ‘deep deletion’ events. Samples with GISTIC values of +1 and −1 were annotated as ‘shallow amplifications’ and ‘shallow deletions’ respectively.

### Determining PPAR, hypoxia and regulatory T cell (Treg) scores

An 18-gene signature of PPAR signaling is developed from putative loss- or gain-of-function genes. Loss-of-function genes were identified from genes that were recurrently deleted and downregulated in tumors. Gain-of-function genes were identified from genes that were recurrently amplified and upregulated in tumors. We calculated 18-gene scores for each patient by taking the average expression of APOA1, PPARA, ACOX2, ANGPTL4, FABP3, PLIN2, AQP7, ACSL1, FABP5, ACADL, PLIN5, PPARG, ACADM, GK, CPT2, SCP2, ACAA1, and PCK2. Hypoxia scores were calculated from the average expression of 52-computationally-derived hypoxia-responsive genes(21). Treg scores were determined from the expression of 31 Treg genes identified f rom t he overlap of four Treg signatures(22–25).

### Differential expression and survival analyses

We have published detailed methods on both types of analyses(26); hence they will not be repeated here. Briefly, differential expression analyses on 74 PPAR pathway genes were performed on tumor and non-tumor samples. Differential expression analyses were also performed between the 4th and 1st quartile patients stratified u sing t he 18-gene signature. Survival analyses were performed using the Kaplan-Meier method coupled with log-rank tests, Cox proportional hazards regression and receiver operating characteristics. For figures 5 and 6, Spearman’s correlation analyses were performed to determine the relationship between PPAR signaling and tumor hypoxia or Treg expression. Patients were separated into four categories based on median PPAR 18-gene scores and hypoxia or Treg scores for Kaplan-Meier and Cox regression analyses. Circular heatmaps in figure 7 were generated based on 18-gene scores in glioma patients ranked in decreasing order from high (purple) to low (yellow). Immune checkpoint genes (PD1, PDL1, PDL2, CTLA4, and CD276) were sorted according to decreasing 18-gene scores for heatmap generation.

**Fig. 1.**
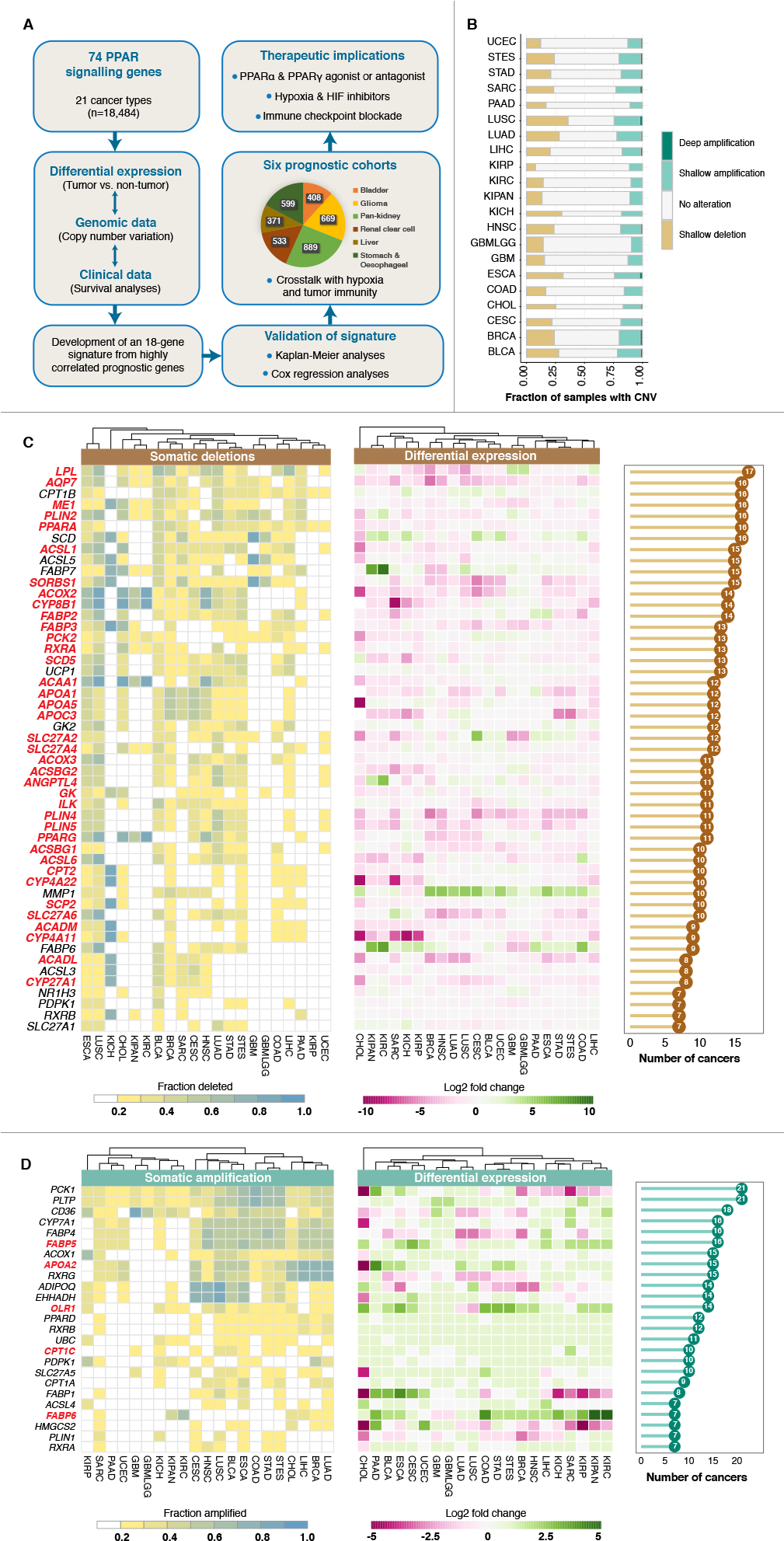
Pan-cancer genomic alterations of PPAR pathway genes. (A) Development of an 18-gene signature consisting of putative loss- and gain-of-function drivers. Somatic copy number alterations and differential expression of 74 PPAR genes are investigated. Prognosis of the signature is validated in six cancer cohorts. Pie slices depict the number of patients within each cohort. Crosstalk between PPAR signaling, tumor hypoxia, and tumor immunity is investigated. (B) Stacked bar chart depicts the fraction of samples within each cancer that contain shallow and deep copy number alterations. The width of the bars is proportional to the number of samples within each cancer type. (C) Somatic losses and differential expression profiles of 51 genes and (D) somatic gains and differential expression profiles of 25 genes that are recurrently deleted or amplified respectively in at least 20% of samples within each cancer in at least 7 cancer types. Heatmaps on the far left show the fraction of samples in which a given gene is deleted or amplified. Heatmaps in the center show differential expression values between tumor and non-tumor samples. Cancer types are ordered using hierarchical clustering with Euclidean distance metric. Genes that are lost and downregulated (38 genes) and genes that are gained and upregulated (5 genes) are highlighted in red. Bar charts on the far right depict the number of cancers harboring at least 20% of samples affected by gains and losses. Refer to Table S2 for cancer abbreviations.

### KEGG enrichment, Gene Ontology (GO) and transcription factor analyses

Differentially expressed genes (DEGs) from 4th vs. 1st quartile patients were used for pathway analyses using GeneCodis(27) and Enrichr(28, 29). DEGs were mapped to KEGG and GO databases to identify significantly enriched pathways. DEGs were mapped to ChEA and ENCODE databases to identify transcription factors involved in regulating the DEGs. All plots were generated using R packages (pheatmap, gg-plot2, and GOplot). The Venn diagram was generated using InteractiVenn(30).

## Results

### Somatic copy number alterations of PPAR pathway genes reveal conserved driver mutations that were recurrently amplified o r deleted

We retrieved 74 genes implicated in PPAR signaling from the KEGG database along with genomic, transcriptomic and clinical data representing 21 major cancer types (n=18,484) from TCGA (Fig. 1A; Table S2). We analyzed somatic copy number alteration (SCNA) events of the 74 genes in these cancer types and observed that lung squamous cell carcinoma (LUSC) and papillary renal cell carcinoma (KIRP) had the highest and lowest fraction of samples with deleted PPAR pathway genes respectively (Fig. 1B). When considering gene amplification, the highest and lowest fraction of samples with amplified genes were observed in esophageal carcinoma (ESCA) and pancreatic adenocarcinoma (PAAD) respectively (Fig. 1B).

To identify genes that exhibited similar patterns of recurrent deletion or amplification a cross c ancers, w e interrogated SCNA profiles that were present in at least 20% of samples per cancer type in at least one-third of cancer types (> 7 cancers). Using these criteria, we identified 51 and 25 genes that were recurrently lost (Fig. 1C) and gained (Fig. 1D) respectively. All 51 genes were found to be deleted in at least 20% of samples in esophageal carcinoma (ESCA) and lung squamous cell carcinoma (LUSC) (Fig. 1C). In contrast, only 2 out of 51 genes (CPT1B and PPARA) were recurrently deleted in papillary renal cell carcinoma (KIRP) (Fig. 1C). Again, esophageal carcinoma (ESCA) emerged with the highest number of recurrently amplified genes (24 out of 25 genes) while only 3 out of 25 genes (PCK1, PLTP, and CD36) were recurrently amplified in glioma (GBMLGG) (Fig. 1D). Further examination into individual genes revealed that LPL was the most deleted gene found in 17 cancer types, followed by AQP7, CPT1B, ME1, PLIN2, PPARA and SCD that were each recurrently deleted in 16 cancer types (Fig. 1C). In contrast, NR1H3, PDPK1, RXRB, and SLC27A1 were some of the least deleted genes found only in 7 cancer types (Fig. 1C). PCK1 and PLTP were both amplified in all 21 cancers, and four other genes (CD36, CYP7A1, FABP4, and FABP5) were found to be amplified in at least 16 cancer types (Fig. 1D).

We reasoned that SCNA events associated with transcript upregulation or downregulation might represent candidate driver genes. Genes that were concomitantly deleted and downregulated could indicate loss-of-function. Moreover, gain-of-function phenotypes can be predicted from genes that were concurrently amplified and overexpressed. When overlaid with mutation data, differential expression analyses performed between tumor and non-tumor samples identified 43 genes that conformed to these criteria. A total of 38 genes were recurrently deleted and downregulated (log2 fold-change < −0.5, P < 0.05) in at least 7 cancer types: LPL, AQP7, ME1, PLIN2, PPARA, ACSL1, SORBS1, ACOX2, CYP8B1, FABP2, FABP3, PCK2, RXRA, SCD5, ACAA1, APOA1, APOA5, APOC3, SLC27A2, SLC27A4, ACOX3, ACSBG2, ANGPTL4, GK, ILK, PLIN4, PLIN5, PPARG, ACSBG1, ACSL6, CPT2, CYP4A22, SCP2, SLC27A6, ACADM, CYP4A11, ACADL, and CYP27A1 (Fig. 1C). Five genes were found to be amplified and upregulated (log2 fold-change > 0.5, P < 0.05) in at least 7 cancer types: FABP5, APOA2, OLR1, CPT1C, and FABP6 (Fig. 1D). These genes were subsequently prioritized as pan-cancer PPAR driver genes.

### Prognostic significance of highly-correlated PPAR driver mutations in six diverse cancer types

Given the widespread patterns of genomic and transcriptomic alterations of PPAR pathway genes, we reasoned that these features would be significantly associated with patient survival outcomes. Employing Cox proportional hazards regression, we examined the prognostic roles of all 43 driver genes identified previously. Except for ACSBG2, all driver genes harbored prognostic information in at least one cancer type (Fig. 2A). Interestingly, the glioma cohort has the highest number of prognostic genes (30 out of 43) (Fig. 2A). We performed Spearman’s correlation analyses on hazard ratios (HR) retrieved from Cox regression analyses to reveal prognostic genes that symbolize pan-cancer significance. We identified 18 highly-correlated genes in this manner and considered them as a pan-cancer PPAR signature: APOA1, PPARA, ACOX2, ANGPTL4, FABP3, PLIN2, AQP7, ACSL1, FABP5, ACADL, PLIN5, PPARG, ACADM, GK, CPT2, SCP2, ACAA1 and PCK2 (Fig. 2B). We next quantified the extent of PPAR signaling by generating a summary score for each patient based on the average expression of the 18 genes. On average, liver (LIHC) and head and neck cancers (HNSC) had the highest
and lowest levels of PPAR activity respectively (Fig. 2C). When employing the 18-gene scores for patient stratification, we observed that levels of PPAR signaling conferred prognostic information in six cancer cohorts (Fig. 2D and 2E). Interestingly, the significance of PPAR driver genes in determining overall survival is cancer-type dependent. Kaplan-Meier analyses on patients within the 1st and 4th survival quartiles demonstrated that elevated PPAR signaling was significantly correlated with poor prognosis in patients with glioma (P<0.0001) and stomach and esophageal cancers (P=0.0078) (Fig. 2D). On the contrary, high expression of signature genes was associated with better survival outcomes in bladder (P=0.031), pan-kidney (consisting of clear cell renal cell, chromophobe renal cell and papillary renal cell carcinoma; P=0.041), clear cell renal cell (P=0.0027) and liver cancer (P=0.0029) cohorts (Fig. 2D). Univariate Cox regression analyses confirmed that patients with high 18-gene scores (4th quartile) had significantly higher death risks in glioma (HR=6.899, P<0.0001) and stomach and esophageal cancers (HR=1.623, P=0.0084) (Table S3). Likewise, in cancers where PPAR signaling is associated with good outcomes, Cox regression analyses confirmed that patients with high 18-gene scores (4th quartile) had improved overall survival rates, which support the hypothesis on the tumor-attenuating effects of PPAR signaling in these cancers: bladder (HR=0.578, P=0.033), pan-kidney (HR=0.687, P=0.032), clear cell renal cell (HR=0.543, P=0.0041) and liver (HR=0.403, P=0.0063) (Table S3).

**Fig. 2.**
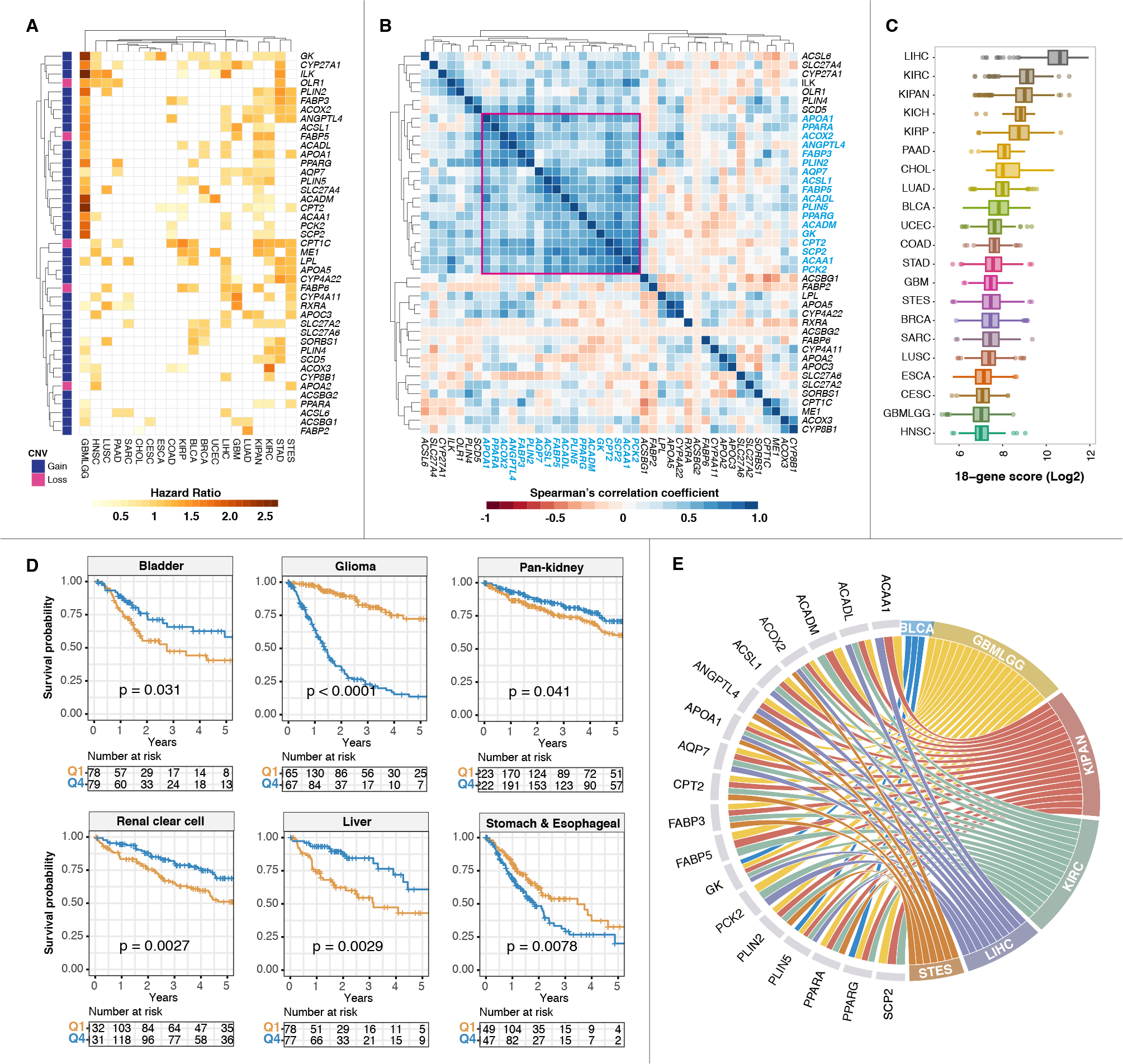
Prognostic significance of PPAR driver genes. (A) Heatmap depicts hazard ratio values obtained from Cox regression analyses on 43 driver candidates across all cancer types. Refer to Table S2 for cancer abbreviations. (B) The heatmap shows Spearman’s correlation coefficient values comparing hazard ratios of the 43 driver genes. Highly correlated prognostic genes are highlighted; gene names marked in blue and correlation coefficients marked with a red box. (C) Box plot represents the distribution of 18-gene scores derived from PPAR signature genes across all cancers. Cancers are ranked from high to low using median scores. (D) Kaplan-Meier analyses of the 18-gene signature confirmed prognosis in six cancer cohorts. Patients are stratified into the 1st and 4th quartiles based on their 18-gene scores. P values are obtained from log-rank tests. (E) Chord diagram shows which of the individual 18 genes harbored prognostic information in the six cohorts shown in (D). For example, GK, PLIN5, and PPARG are individually prognostic in bladder cancer. BLCA = bladder, GBMLGG = glioma, KIPAN = pan-kidney, KIRC = clear cell renal cell, LIHC = liver and STES = stomach and esophageal.

### The 18-gene signature is an independent predictor of survival

The tumor, node, and metastasis (TNM) staging system is frequently used in cancer prognostication. To determine whether the signature was confounded by TNM staging, we performed multivariate Cox regression analyses and found that the PPAR signature remained an independent predictor of overall survival: bladder (HR=0.692, P=0.031), pan-kidney (HR=0.625, P=0.012), clear cell renal cell (HR=0.604, P=0.018), liver (HR=0.492, P=0.035) and stomach and esophageal (HR=1.509, P=0.033) (Table S3). Since the signature was independent of TNM stage, we predict that it could be used to improve the sensitivity and specificity of TNM staging. We employed receiver operating characteristic (ROC) analyses to compare the predictive performance of the signature with a combined model that unites the signature with TNM staging. ROC analyses revealed that the combined model (signature + TNM staging) had higher area under the curve (AUC) values: bladder (0.722 vs. 0.697), pan-kidney (0.811 vs. 0.720), clear cell renal cell (0.813 vs. 0.722), liver (0.745 vs. 0.679) and stomach and esophageal (0.667 vs. 0.601) (Fig. 3B). Moreover, Kaplan-Meier analyses and log-rank tests revealed that the signature offered an additional resolution to further separate similarly-staged tumors: bladder (P<0.0001), pan-kidney (P<0.0001), clear cell renal cell (P<0.0001), liver (P<0.0001) and stomach and esophageal (P<0.0001) (Fig. 3A).

Hyperactivation of PPAR signaling is associated with adverse survival outcomes in glioma patients. Glioma samples are categorized into four histological subtypes: low-grade gliomas (astrocytoma, oligoastrocytoma, and oligodendroglioma) and grade IV glioblastoma multiforme. Kaplan-Meier analyses confirmed that elevated PPAR signaling was indeed associated with poor survival outcomes in astrocytoma (P=0.0011), oligoastrocytoma (P=0.016) and glioblastoma multiforme (P=0.00014) (Fig. 3A). These observations were independently confirmed b y C ox r egression analyses where patients within the 4th quartile had higher mortality rates: astrocytoma (HR=3.204, P=0.0019), oligoastrocytoma (HR=4.232, P=0.027) and glioblastoma multiforme (HR=2.699, P=0.00023) (Table S3). ROC analyses revealed that the predictive performance of the signature was the best in astrocytoma (AUC=0.844), followed by oligoastrocytoma (AUC=0.833) and glioblastoma multiforme (AUC=0.729) (Fig. 3B). The signature also performed well when all histological subtypes (including oligodendroglioma) were considered as a group (AUC=0.832) (Fig. 3B).

### The 18-gene signature reveals aberration in oncogenic pathways converging on similar downstream targets associated with deranged metabolism

Since aberration in PPAR signaling is linked to clinical outcomes (Fig. 2 and 3), we hypothesized that the function of driver genes in these cancers might converge on similar downstream targets. To identify oncogenic targets associated with PPAR signaling, we performed differential expression analyses between patients separated into 4th and 1st quartiles based on their 18-gene scores. The highest number of differentially expressed genes (DEGs; −1.5 > log2 fold change > 1.5; P<0.01) was observed in glioma (2,240 genes), followed by liver (1,578 genes), bladder (1,374 genes), stomach and esophageal (1,323 genes), pan-kidney (897 genes) and clear cell renal cell (721 genes) (Fig. 4A; Table S4). Remarkably, we observed overlaps in downstream transcriptional targets resulting from aberrant PPAR signaling in these cancers despite their distinct pathologies; 23 genes were found to be dysregulated in at least five cohorts, 249 genes in at least four cohorts and 739 genes in at least three cohorts (Fig. 4A; Table S4). Together, these implied that PPAR signaling plays a conserved role in driving oncogenic progression. We performed Gene Ontology (GO) and KEGG enrichment analyses to further assess the functional significance of the DEGs (Table S4). All six cancer cohorts exhibited very similar patterns of enriched biological processes and oncogenic pathways (Fig. 4B and 4C). Biological functions associated with metabolism, cell adhesion, and cell-to-cell signaling were among some of the most enriched processes (Fig. 4B). Moreover, KEGG analyses showed that numerous metabolic-related pathways were dysregulated in patients with altered PPAR signaling (Fig. 4C), which provided an independent confirmation of the functional significance of the PPAR pathway in lipid and fatty acid metabolism. Together, these results suggest that altered PPAR signaling may cause metabolic reprogramming of tumor cells to impact patient survival directly. To determine which transcription factors (TFs) were upstream regulators of the DEGs, we mapped the DEGs to ENCODE and ChEA databases using the Enrichr tool. Interestingly, DEGs from all six cohorts were enriched for binding targets of SUZ12 and EZH2, which are important regulators of cancer stem cells (CSCs) (Fig. 4D). Enrichments of other TFs implicated in CSC maintenance, RE1 Silencing Transcription Factor (REST) and SMAD4, were also observed, implying the underappreciated connection between PPAR signaling and self-renewal mechanisms (Fig. 4D).

### Tumor hypoxia worsen survival outcomes in patients with impaired PPAR signaling

Hypoxia is a universal feature in almost all solid malignancies due to the formation of aberrant tumor microvasculature(31–34). As an adaptation strategy to hypoxic environments, tumor cells need to reprogram their metabolic requirements to survive(35). Since PPARs are critical metabolic regulators, we predict that tumor hypoxia would influence the behavior of PPAR pathway genes and consequently patient prognosis. To evaluate the contribution of tumor hypoxia on PPAR signaling, we assessed the levels of hypoxia in each patient using a hypoxia gene signature where hypoxia scores were calculated from the mean expression values of 52 hypoxia-responsive genes(21). PPAR scores were significantly positively correlated with hypoxia scores in glioma (rho=0.64, P<0.0001) and pan-kidney (rho=0.24, P<0.0001) cohorts (Fig. 5A). However, this trend was reversed in liver (rho= −0.47, P<0.0001) and stomach and esophageal cancers (rho= −0.20, P<0.0001) (Fig. 5A). Patients were grouped into four categories based on their median PPAR and hypoxia scores for survival analyses. Intriguingly, the joint relationship between hypoxia and PPAR signaling allowed further delineation of risk groups that influenced survival outcomes: glioma (full cohort; P<0.0001), astrocytoma (P<0.0001), oligoastrocytoma (P=0.001), pan-kidney (P<0.0001), liver (P=0.0007) and stomach and esophageal (P=0.039) (Fig. 5B). According to analyses in the previous section, high 18-gene PPAR scores can either be associated with poor or good outcomes depending on the cancer type (Fig. 2). In cancers where a high PPAR score is associated with poor outcomes, a high hypoxia score would further exacerbate disease phenotypes in these patients: glioma (full cohort; HR=7.938, P<0.0001), astrocytoma (HR=7.380, P<0.0001), oligoastrocytoma (HR=14.179, P=0.011) and stomach and esophageal (HR=1.748, P=0.0058) (Fig. 5C). On the other hand, high PPAR score is a good prognostic factor in renal and liver cancers. Hence, patients with both low PPAR and high hypoxia scores would have the highest mortality rates: pan-kidney (HR=3.187, P<0.0001) and liver (HR=2.849, P=0.00014) (Fig. 5C). These results suggest a model whereby PPAR signaling may influence transcriptional targets of hypoxia.

**Fig. 3.**
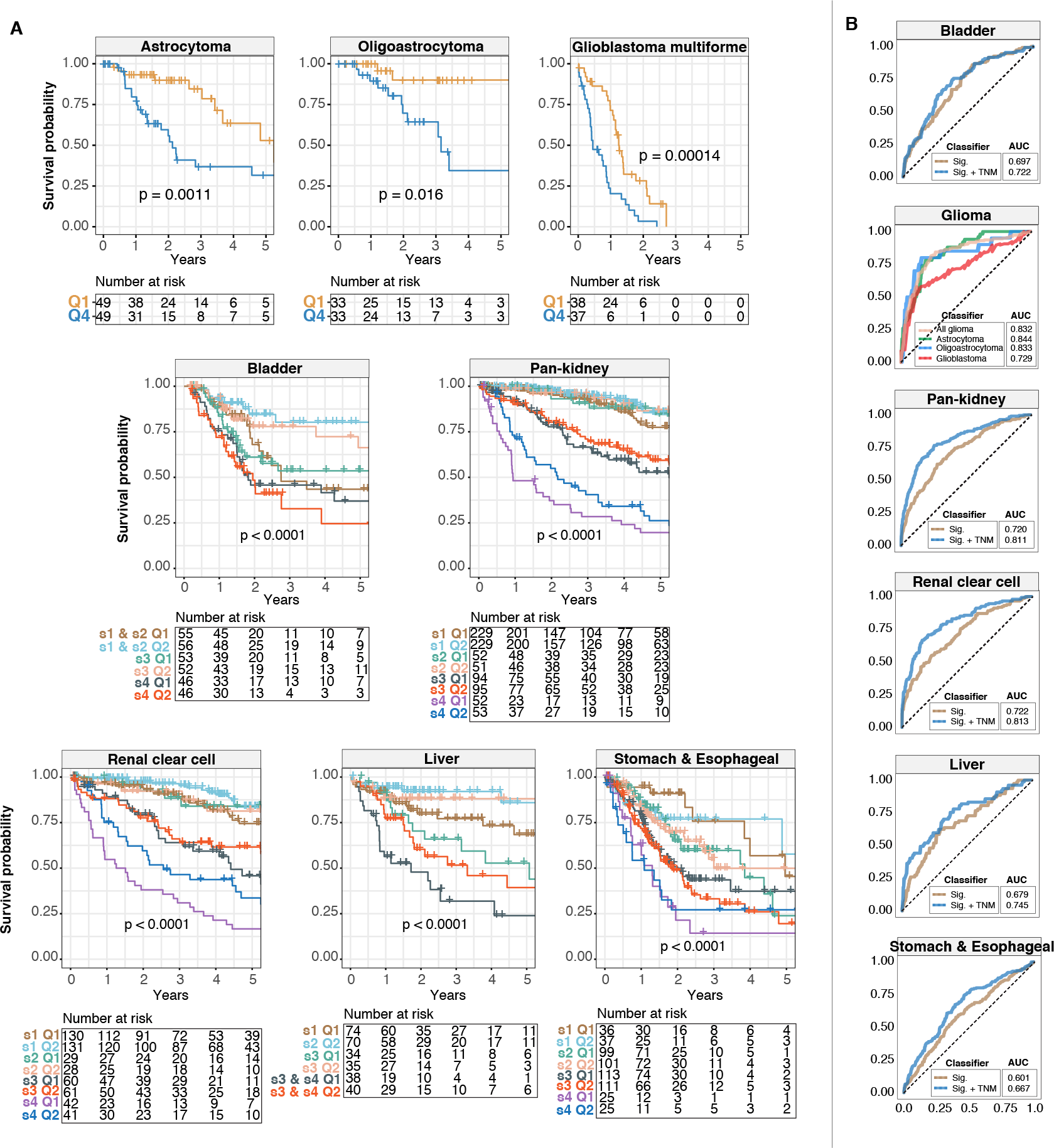
The 18-gene signature is independent of tumor stage. (A) Kaplan-Meier analyses of patients stratified by tumor stage or in the case of glioma by histological subtype, and the 18-gene signature. For histological subtypes of glioma, log-rank tests are used to compare patients within the 1st and 4th survival quartiles. For the other cancers, patients are first stratified according to tumor, node, and metastasis (TNM) staging followed by median-stratification into low- and high-score groups using the 18-gene signature. P values are obtained from log-rank tests. (B) Predictive performance of the signature is determined using receiver operating characteristic analyses. ROC curves generated using the signature are compared to those generated using a combined model uniting the signature and TNM staging. For glioma patients, the area under the curves (AUCs) for astrocytoma, oligoastrocytoma, glioblastoma multiforme and all glioma samples as a whole are shown.

**Fig. 4.**
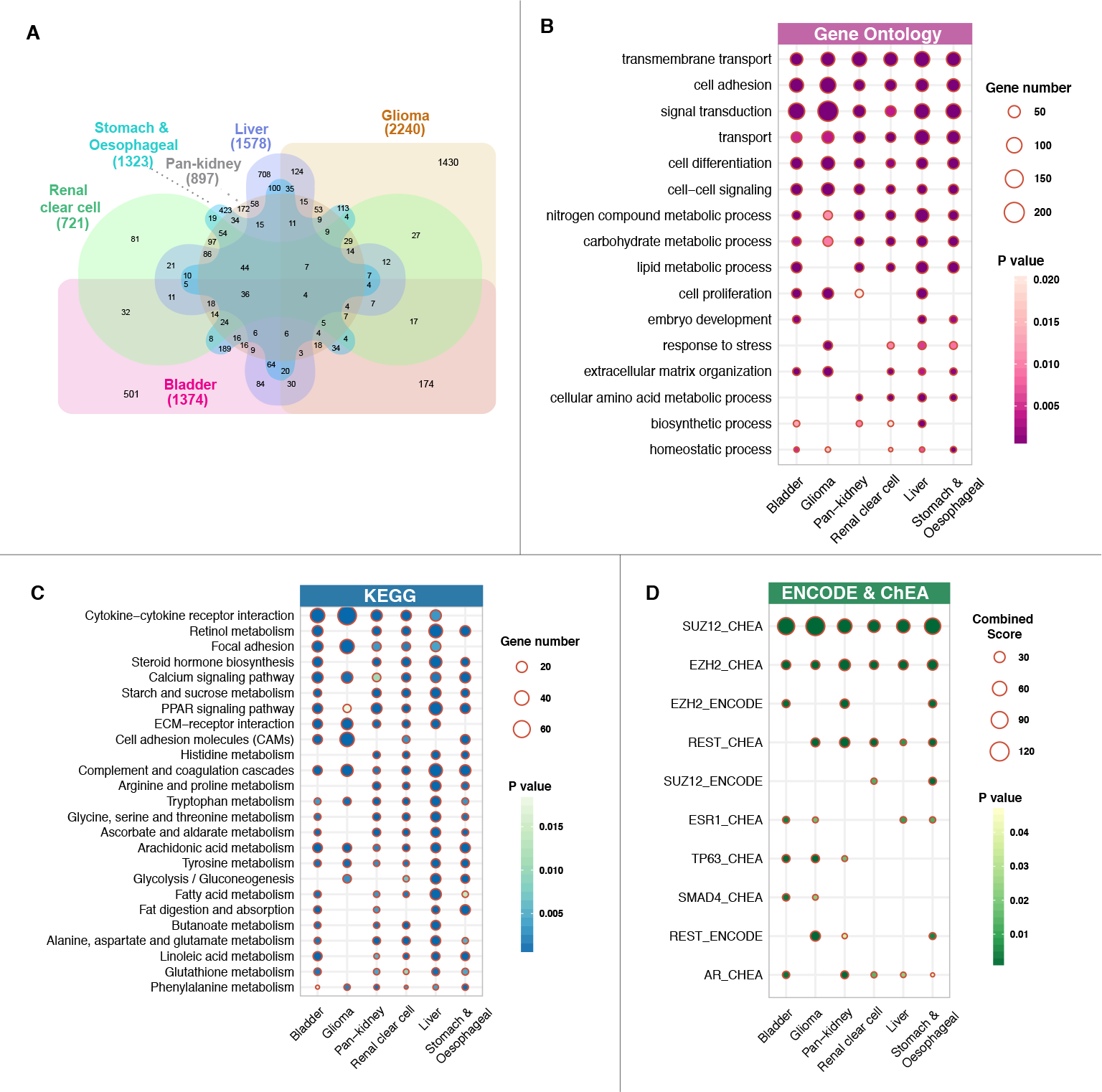
Dysregulated PPAR signaling converges on transcriptional targets implicated in diverse metabolic processes. Differential expression analyses are performed between 4th and 1st quartile patients stratified by the 18-gene signature in six cancer cohorts. (A) The Venn diagram shows the number of overlapping differentially expressed genes (DEGs) in the six cohorts. DEG numbers are shown in parentheses. Functional enrichment analyses are performed by mapping DEGs to the (B) Gene Ontology, (C) KEGG and (D) ENCODE and ChEA databases. Enriched transcription factor binding associated with DEGs are shown in (D).

### Immuno-oncogenic properties of PPAR driver genes in patients with glioma

It was reported recently that focal amplification a nd over-expression of PPARγ inhibits the secretion of inflammatory cytokines and reduces cytotoxic CD8+ T-cell infiltration(36). PPARγ and RXRA activities promote resistance to immune checkpoint blockade to create an environment that favors tumor growth(36). In order to evaluate the role of PPAR signaling in modulating tumor immunity, we first needed to identify genes that are associated with immunosuppression. We retrieved four regulatory T cell (Treg) gene signatures(22–25) and looked for genes that were common in all four signatures. We identified 31 genes that were present in all four signatures to yield a more representative Treg gene set that is not specific to single tumor type. We calculated Treg scores for each patient based on the average expression of the 31 genes. A strong positive correlation between Treg and PPAR 18-gene scores (rho=0.65, P<0.0001) was observed in glioma patients, suggesting that tumor cells with elevated PPAR signaling were hypoimmunogenic (Fig. 6A). Patients were stratified into four categories based on median PPAR and Treg scores for survival analyses. Intriguingly, high expression of Treg genes would further promote disease aggression in glioma tumors with hyperactive PPAR signaling (HR=7.356, P<0.0001) (Fig. 6B and 6C). This observation was repeated in glioma histological subtypes: astrocytoma (HR=3.699, P<0.0001) and oligoastrocytoma (HR=3.227, P=0.038) (Fig. 6B and 6C). Taken together, metabolic reprogramming of the tumor microenvironment through PPAR signaling may influence anti-tumor immunity and consequently immunotherapeutic outcomes in clinical settings. Indeed, we observed that PPAR signaling is significantly positively correlated with the expression profiles of canonical immune checkpoint markers, suggesting that pharmacological inhibition of PPAR could reinvigorate immunosurveillance in glioma patients; PD1 (rho=0.41, P<0.0001), PDL1 (rho=0.49, P<0.0001), PDL2 (rho=0.62, P<0.0001), CTLA4 (rho=0.38, P<0.0001) and CD276 (rho=0.36, P<0.0001) (Fig. 7C).

**Fig. 5.**
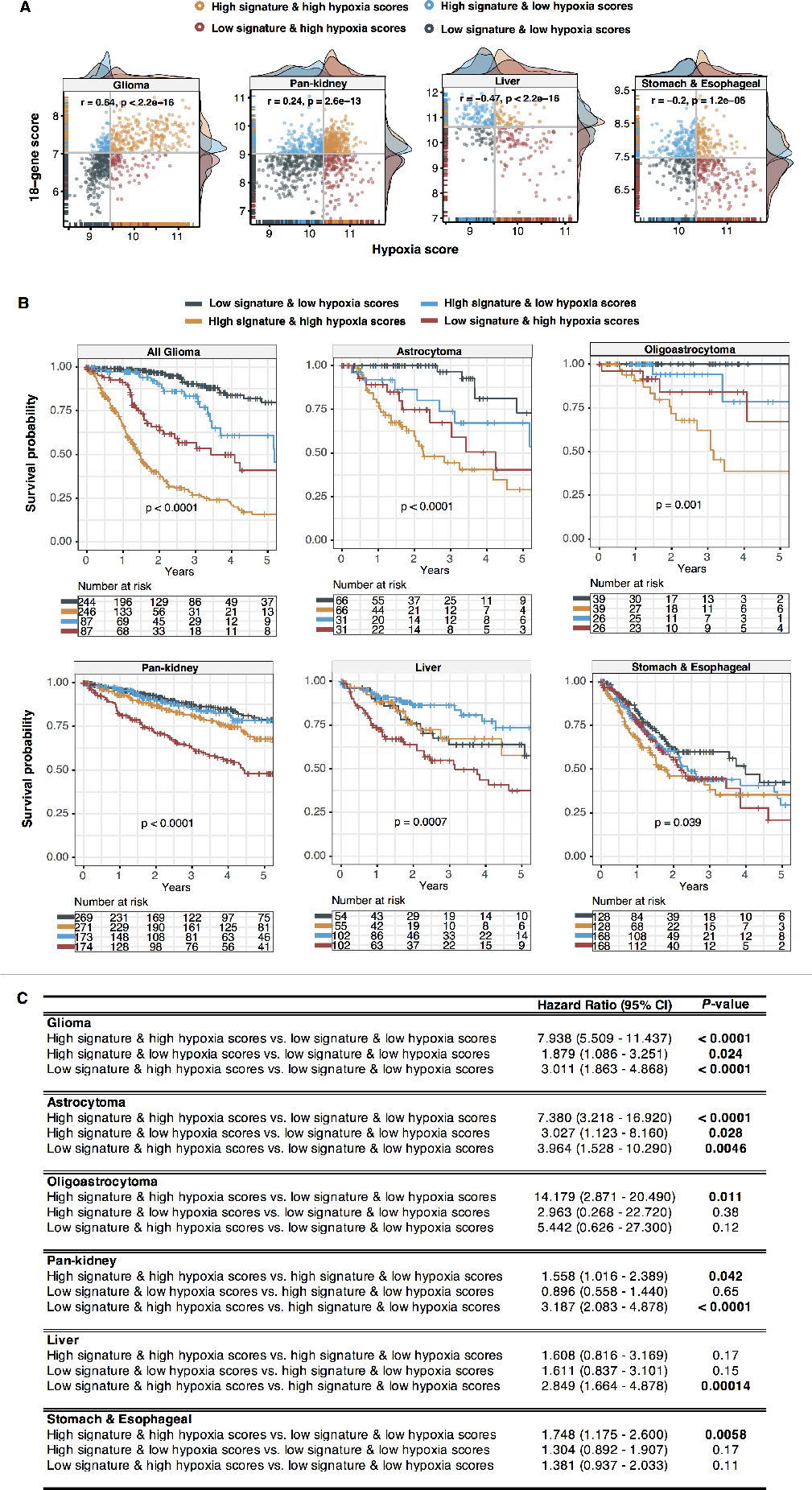
Prognostic relevance of the crosstalk between PPAR signaling and tumor hypoxia. (A) Scatter plots show significant positive or negative correlations between 18-gene and hypoxia scores. Patients are grouped into four categories based on median 18-gene and hypoxia scores. Density plots at the x- and y-axes show the distribution of 18-gene and hypoxia scores. (B) Kaplan-Meier analyses are performed on the four patient groups to determine the ability of the combined PPAR-hypoxia model in determining overall survival in multiple cancers including glioma histological subtypes. P values are obtained from log-rank tests. (C) Table inset shows univariate Cox proportional hazards analyses of the relationship between PPAR signaling and hypoxia. Significant P values are highlighted in bold. CI = confidence interval.

## Discussion

In an integrated analysis involving multidimensional datasets from TCGA, we examined molecular characteristics of the PPAR pathway comprising of 74 genes across 18,484 patients representing 21 cancer types. We started by cataloging driver genes with significant SCNA patterns linked to transcriptional dysregulation. We uncovered tissue-specific and universal mutation profiles that resulted in gene upregulation or silencing, which highlight novel oncogenic mechanisms of PPAR signaling that has been previously underappreciated. We identified a set of highly correlated genes (18-gene signature) that were consistently associated with survival outcomes across six cancer cohorts. In-depth analysis of differential PPAR signaling uncovered transcriptional perturbations of numerous signal-transduction pathways that were conserved among cancer types. Pathways associated with metabolic processes were among the most enriched ontologies; an observation that independently validates the role of PPARs in lipid transport, fatty acid oxidation, fatty acid catabolism and crosstalk with other lipogenic pathways(37).

Studies have demonstrated opposing effects of PPARs, whereby they could either promote or inhibit tumor growth. PPARδ, allows breast cancer cells to persist in severe metabolic environments; its expression level is negatively correlated with survival outcomes and increased metastatic ability of tumor cells in mice(38). In pancreatic cancer, PPARδ represents a hub gene of a transcriptome-derived angiogenic network(39). Suppression of PPARδ inhibits tumor growth and angiogenesis since its expression is positively correlated with tumor aggression, recurrent and metastasis(39). ANGPTL4 is a well-established PPARδ target gene. It promotes cell migration in colon cancer cells(40), lung metastasis in breast cancer(41) and venous invasion in colorectal and gastric cancers(42, 43). Moreover, inhibition of ANGPTL4, a downstream target of PPARδ, abrogates its pro-oncogenic effects and suppresses breast cancer cell invasion(44). In our study, we identified ANGPTL4 as one of the PPAR pathway driver genes. We found that the expression level of ANGPTL4 is positively correlated with poor survival outcomes in glioma, stomach and esophageal, lung, colon and renal cancers and good survival outcomes in liver cancer (Fig. 2A; Fig. 7A). PPARs are highly pleiotropic since they also possess anti-tumor functions. PPARγ induces apoptosis and inhibits tumor growth in colon cancer cells(45). PPARγ ligands impair gastric cancer cell proliferation in a dose-dependent manner due to the upregulation of p53 and downregulation of cyclin E1 and tumor burden in mice was reduced after treatment with a PPARγ agonist rosiglitazone(46). Moreover, over-expression of PPARγ inhibits the metastatic potential of gastric cancer cells through attenuation of Wnt/β-Catenin signaling and TERT expression(47). Our results demonstrate that elevated expression of PPARγ is associated with better survival outcomes in renal and bladder cancers, but poor survival outcomes in glioma (Fig. 2A; Fig. 7A). These striking cancer type-dependent patterns further reinforce the benefit of our study, which investigates PPAR pathway alterations in a larger pan-cancer context to expose diverse molecular complexities and cancer-specific vulnerabilities.

**Fig. 6.**
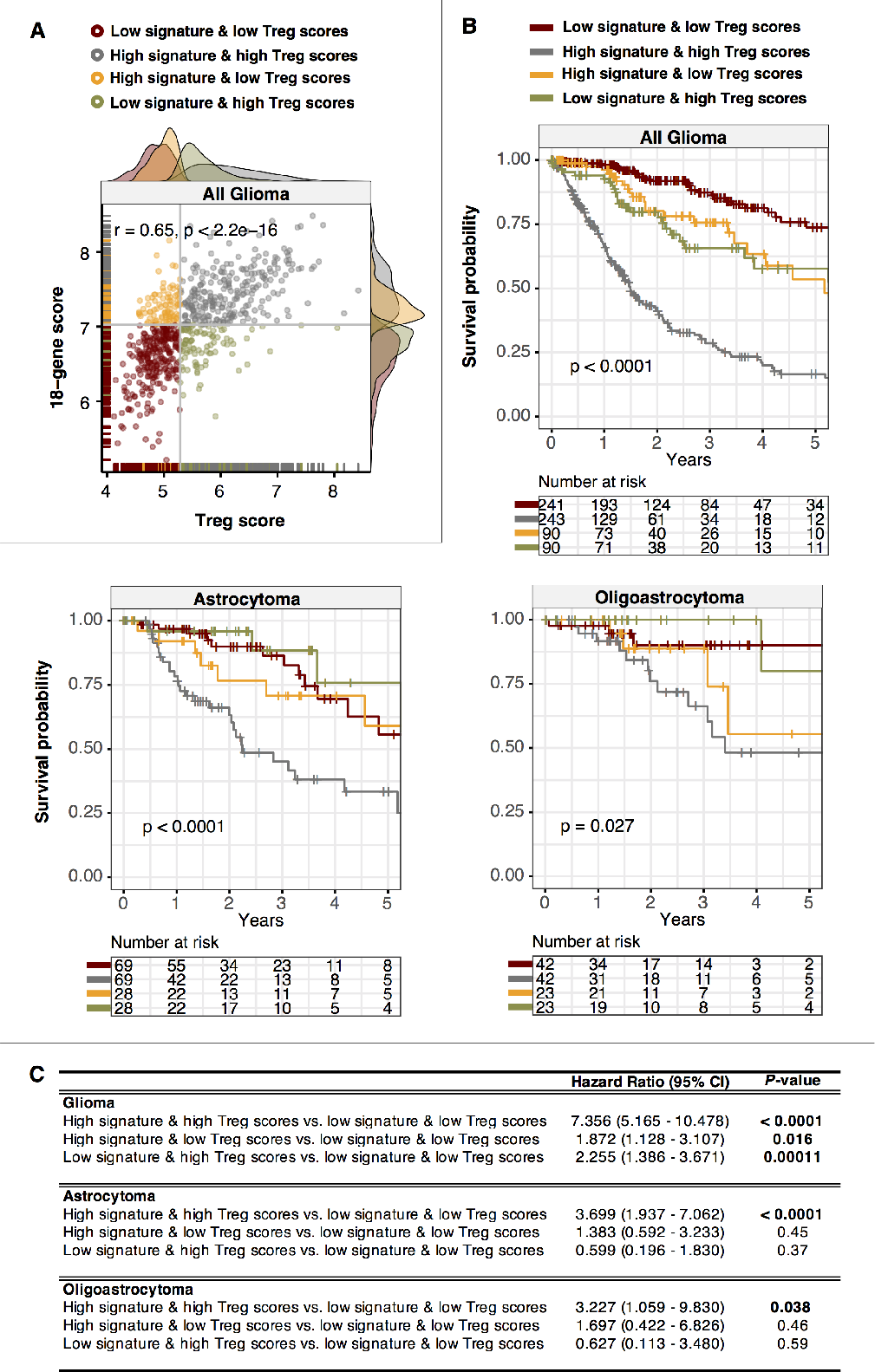
PPAR signaling is associated with immunosuppressive phenotypes. (A) Scatter plot depicts a significant positive correlation between 18-gene and regulatory T cell (Treg) scores in glioma patients. Patients are grouped into four categories based on median 18-gene and Treg scores. Density plots at the x- and y-axes show the distribution of 18-gene and Treg scores. (B) Kaplan-Meier analyses are performed on the four patient groups to determine the ability of the combined PPAR-Treg model in determining overall survival in glioma histological subtypes. P values are obtained from log-rank tests. (C) Table inset shows univariate Cox proportional hazards analyses of the relationship between PPAR signaling and tumor immunity in glioma. Significant P values are highlighted in bold. CI = confidence interval.

Metabolic signals in the tumor microenvironment may regulate the behavior of immune cells and affect patient response to immunotherapies. Immune cells possess unique metabolic qualities and bioenergetic requirements(48). Thus, alteration of the microenvironmental metabolic landscape could have a direct impact on anti- or pro-tumor effects(48). As lipid-sensitive nuclear receptors, it is therefore not surprising that PPARs could modulate metabolic homeostasis of immune cells. Indeed, multiple studies have shed light on the defining roles of PPARs in regulating innate and adaptive immunity responses(49). PPARγ activation in macrophages stimulates lung cancer metastasis(50), while PPARδ activation promotes apoptotic cell clearance(51). PPARδ activation in myeloid cells promotes tumor invasion through the activation of interleukin 10(52). Also, dysregulated lipid metabolism and accumulation of fatty acids in non-alcoholic fatty liver disease patients lead to increased oxidative damage, the loss of CD4+ T helper cells and impaired anti-tumor immunity in liver cancer(53). A majority of patients with muscle-invasive bladder cancer (MIBC) do not respond to immunotherapy. Korpal et al. found that immune evasion in MIBC patients is caused by PPARγ/RXRA overexpression, which inhibits inflammatory cytokine release(36). Cytokine expression is restored through pharmacological inhibition of PPARγ(36). Moreover, activation of LXRs (another member of the nuclear receptor superfamily involved in lipid homeostasis) inhibits neutrophil and dendritic cell migration into tumors contributing to tumor immunotolerance(54). Our study may help prioritize PPAR candidates for future functional studies to pave the way for successful immunotherapeutic approaches.

Oxygen deprivation or hypoxia within the tumor microenvironment contributes to additional metabolic stress where hypoxia could drive the reprogramming of metabolic genes as an adaptation strategy for tumor survival(35). We found that PPAR signaling can either be positively or negatively correlated with tumor hypoxia depending on cellular context (Fig. 7B). PPARγ expression and activity levels are downregulated under hypoxic conditions in human pulmonary artery smooth muscle cells leading to pulmonary hypertension(55). PPARs could, in turn, regulate HIF-1a signaling in breast and ovarian cancer cell lines; treatment of these cells with a PPAR agonist before hypoxia incubation promotes HIF-1a degradation and suppresses VEGF secretion resulting in anti-tumor effects(56). Curiously, another study reported that hypoxia elicited the downregulation of PPARα in intestinal epithelial cells(57) and since PPARα inhibits HIF-1a signaling, the absence of PPARα may be crucial for cells to mount an appropriate hypoxic response. We recently demonstrated that hypoxia also promotes the expression of Tregs and contributes to the impairment of anti-tumor surveillance to directly impact patient prognosis(26). Hypoxia modulates angiogenic processes through regulating the function of immune cells. In natural killer cells, HIF-1a accumulation promotes tumor growth by inhibiting non-productive angiogenesis(58). Hypoxia also promotes the angiogenic and immunosuppressive features of tumor-associated macrophages leading to tumor metastasis(59). Immunosuppressive myeloid-derived suppressor cells are recruited to hypoxic liver tumors and are blocked by HIF inhibitors to reestablish immune surveillance(60). Further interrogation of the metabolism-hypoxia-immunity axis would provide a framework for combination therapeutic initiatives to simultaneously target these pathways.

**Fig. 7.**
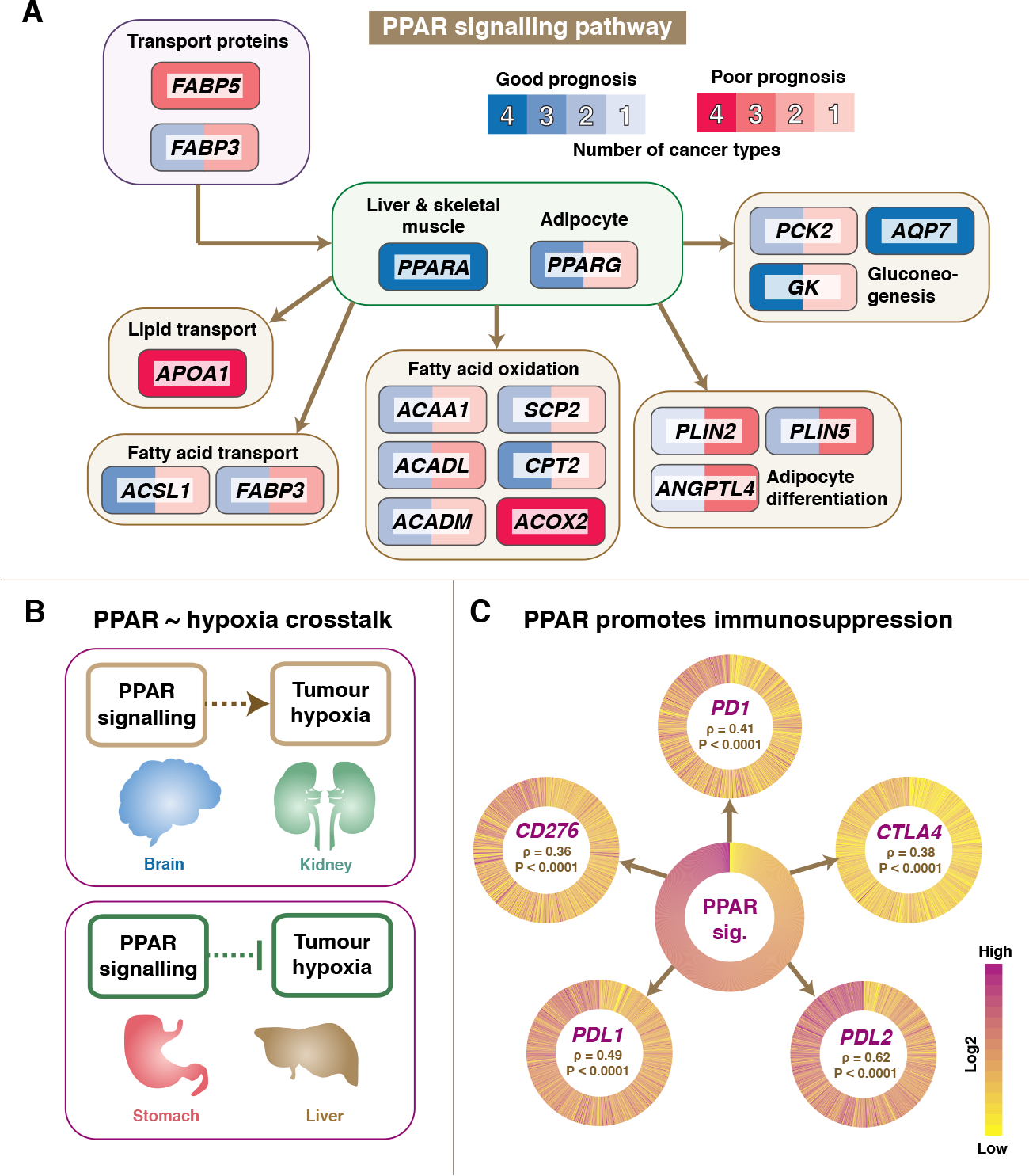
Pan-cancer model of PPAR dysregulation and proposed crosstalk with tumor hypoxia immunity. (A) The pathway diagram represents the relationship between each of the 18 signature genes. Genes that confer good prognosis are colored in blue while genes that confer poor prognosis are colored in red. Color intensities represent the number of cancer cohorts in which a given gene is prognostic. Only six cohorts, as shown in figures 2D, 3 and 4, are considered in this diagram. (B) As determined in figure 5, PPAR signaling is positively or negatively correlated with tumor hypoxia depending on cellular context. (C) PPAR signaling promotes immune evasion in glioma. The circular heatmap in the center represents 18-gene scores in glioma sorted in descending order with each spoke representing an individual patient. Circular heatmaps of five immune checkpoint genes are plotted with patients sorted in descending order of 18-gene scores. Spearman’s correlation coefficients between 18-gene scores and immune checkpoint gene expression values are shown in the center of the heatmap.

## Conclusion

In summary, we present a comprehensive catalog of genetic variants associated with PPAR signaling in over 18,000 tumor samples, including clinically actionable driver genes, which may serve as important foundations for understanding the wide-ranging effects of PPARs in tumorigenesis. Our study reveals candidate driver genes that may be used for personalized cancer treatment utilizing small molecule inhibitors or agonists targeting the PPAR pathway. Furthermore, the 18-gene signature could be used for patient stratification in cancer therapy involving hypoxia inhibitors and/or immune checkpoint blockade.

